# Speaking in the brain: The interaction between words and syntax in producing sentences

**DOI:** 10.1101/696310

**Authors:** Atsuko Takashima, Agnieszka Konopka, Antje Meyer, Peter Hagoort, Kirsten Weber

## Abstract

This neuroimaging study investigated the neural infrastructure of sentence-level language production. We compared brain activation patterns, as measured with BOLD-fMRI, during production of sentences which differed in verb argument structures (intransitives, transitives, ditransitives) and the lexical status of the verb (known verbs or pseudo-verbs). An example for the type of sentence to be produced started a mini-block of six sentences with the same structure. For each trial, participants were first given the (pseudo-)verb followed by three geometric shapes to serve as verb arguments in the sentences. Production of sentences with known verbs yielded greater activation compared to those with pseudo-verbs in the core language network of left inferior frontal gyrus, the left posterior middle temporal gyrus, and a more posterior middle temporal region extending into the angular gyrus (LpMTG/AG), analogous to effects observed in language comprehension. Increasing the number of verb arguments led to greater activation in an overlapping left pMTG/AG area, particularly for known verbs, as well as in the bilateral precuneus. Thus, producing sentences with more complex structures using existing verbs lead to increased activation in the language network, suggesting some reliance on memory retrieval of stored lexical-syntactic information during sentence production. This study thus provides evidence from sentence-level language production in line with functional models of the language network that have so far been mainly based on single word production, comprehension and processing in aphasia.

---

Before initiating production, speakers must select relevant words from their mental dictionary and assemble these words together in a suitable syntactic structure in order to produce an utterance that conveys the desired message in a grammatically correct form. Based on behavioural data, psycholinguists have proposed two accounts of lexical-structural processing during production: a lexicalist account where the retrieval of individual words may activate syntactic information tied to those words (Bock, 1982; Pickering & Branigan, 1998; Tomasello, 2000) and an abstract structural account where the selection and generation of a suitable structure is driven by message-level constraints and thus can proceed independently of lexical retrieval (Bock, 1986; Chang, Dell, & Bock, 2006; Frazier, 1987). Speakers are able to form grammatically plausible sentences with novel words that do not have entries in the mental lexicon, suggesting that they must have acquired sentence structure templates in the course of language learning that are at least partially abstract and not bound to specific words. This debate has motivated behavioural investigations into the time-course of structural and lexical processing during sentence production using a range of experimental paradigms (such as object naming, spontaneous event description, structural priming; e.g., Allum & Wheeldon, 2007; Konopka, 2012; Smith & Wheeldon, 1999). Broadly speaking, there is now evidence that utterance production relies on abstract syntactic structures but also that the accessibility of lexical information can influence the process of structural assembly.

Here we investigate the neural underpinnings of sentence production with different syntactic structures. Neural models of language processing propose different functional roles for the different subcomponents of this language network. In the MUC model for example, the posterior temporal cortices subserve memory (M), the inferior frontal cortex including Broca’s area and adjacent structures supports unification (U), and the dorsal lateral and medial prefrontal cortex (including the anterior cingulate cortex) have been related to control (C) processes (Hagoort, 2005, 2013). This model (Hagoort, 2005, 2013) and a related computational implementation (Vosse & Kempen, 2000) propose that each entry in the mental lexicon is linked to its structural information. For example, verb argument structure is verb-bound and the syntactic role of these arguments must be defined for each verb (e.g., an intransitive verb will require only a subject, whereas a ditransitive verb will require a subject, a direct object and an indirect object). This view is compatible with lexicalist accounts in (psycho)linguistics (e.g., (Bresnan, Asudeh, Toivonen, & Wechsler, 2015; Jackendoff, 2002; Pickering & Branigan, 1998) in which lexical items co-determine the temporal dynamics and the outcome of syntactic encoding processes. A mental lexicon in the posterior temporal cortex that drives lexical-syntactic processing, however, does not exclude the possibility that more abstract structural information can also be processed or unified by other regions involved in syntactic processing, such as the left inferior frontal gyrus.

Existing neural models of language processing are based mainly on data from comprehension studies. FMRI studies on language production studies have been scarce due to presence of movement artefacts during speaking. There are, however, ways to tackle this problem by including a motion predictor regressor in the model for functional magnetic resonance imaging (fMRI) data. Here, we focus on processing of sentence structure. Earlier meta-analyses of neuroimaging studies on language processing in healthy participants have identified the left frontal and temporal cortices as important structures for syntactic processing (Indefrey & Levelt, 2004; Price, 2012). In a PET study, an increase in left inferior frontal gyrus (LIFG) activation was observed when participants produced full sentences compared to when they produced single words (Indefrey et al., 2001). In an fMRI study, an increase in activation was found in the LIFG when participants generated sentences cued by visual scenes or a scrambled word orders (e.g., cue: “Throw ball child”, response: *“The child throws the ball”),* compared to when the participants read out the words on the screen in a scrambled order (e.g., “throw ball child”) or read the complete sentence presented on the screen (e.g., “The child throws the ball”). This suggests a functional role of the LIFG in syntactic encoding.

More compelling evidence for the organisation of the language network comes from priming studies showing that this left-dominant network of the left inferior frontal, middle and superior temporal cortex involved in sentence production largely overlaps with the comprehension network. Repeating syntactic structures, semantic contents or lexical items lead to repetition suppression, i.e., a decrease in the neuronal activation of the regions involved in processing the repeated features (reviewed in Barron, Garvert, & Behrens, 2016; Segaert, Weber, de Lange, Petersson, & Hagoort, 2013). Repetition of syntactic structures results in suppression of activation in the left inferior frontal, precentral and posterior temporal regions, repetition of semantic information in the bilateral posterior temporal and precuneus, and repetition of words in the anterior and posterior temporal and anterior inferior frontal regions and precuneus. These repetition suppression effects are not only observed within modality but also between comprehension and production (Menenti, Gierhan, Segaert, & Hagoort, 2011; Segaert, Menenti, Weber, Petersson, & Hagoort, 2012). However, as neuroimaging studies have mostly considered a very limited set of sentence structures (mostly transitives; Haller, Radue, Erb, Grodd, & Kircher, 2005; Indefrey et al., 2001; Menenti et al., 2011; Segaert et al., 2012), we know much less about the production of sentences with different levels of complexity.

Regarding complex structure sentence processing, studies on patients with aphasia shed some light on the brain areas involved. A growing number of observations in aphasic studies found the left inferior frontal cortex to be critical in syntactic processing (see recent review by Tyler et al., 2011), although this view is also contested (W. Matchin & Hickok, in press). For verbs, syntactic complexity is related strongly to argument number. Intransitive, transitive, and ditransitive verbs are bound to different numbers of argument slots (intransitive – 0 slots: “*The girl sleeps*.”, transitive – 1 slot: “*The girls kicks <the ball*>.”, ditransitive – 2 slots: *“The girl gives <the book> < to the boy*>.”). This sensitivity is lacking in the individuals with Wernicke’s aphasia whose lesions are found in the posterior perisylvian regions. A recent study training patients with agrammatic aphasia to produce ditransitive sentences from pictures of simple events (e.g., *“The boy is giving the flowers to the woman*”) showed an improvement in naming of trained three-argument verbs in isolation, as well as production of these verbs in full sentences. These patients also showed an increased activity pattern in the angular, supramarginal and/or superior posterior temporal gyri during an action verb video naming task (Thompson, Riley, den Ouden, Meltzer-Asscher, & Lukic, 2013).

Lesion studies are invaluable in understanding the neural correlates of language production, but the results may be affected by compensatory mechanisms and altered neural responses due to regional deficits. Thus, it is also important to investigate these effects in a healthy population. Neuroimaging studies have found a gradient of activation with an increase in number of verb arguments in the posterior temporal and inferior parietal parts of the language network. The studies, however, have mainly focused on verbs comprehended or produced in isolation. That is, the stimuli were verbs with different types of verb argument structure (intransitive/transitive/ditransitive) presented outside of a sentence context (e.g., Malyutina & den Ouden, 2017, Exp 2; Thompson et al., 2007). Other studies tested grammaticality, or well-formedness judgements for sentences in comprehension paradigms (Ben-Shachar, Palti, & Grodzinsky, 2004; Malyutina & den Ouden, 2017, Exp 1). Nevertheless, all these studies found activation differences in the posterior perisylvian area. These findings suggest that verbs may be stored in the mental lexicon together with their syntactic information and are therefore consistent with lexicalist accounts of structural processing (Jackendoff, 2002; Pickering & Branigan, 1998).

However, as these studies did not examine *sentence-level* production, they cannot speak to the way these verbs are retrieved and used in more complex contexts, such as during the generation of full sentences. Moreover, a problem in the previous single verb production studies is that the images or videos used to cue the production of verbs with different argument structures often contained different numbers of depicted objects (e.g., 1 vs. 2; den Ouden, Fix, Parrish, & Thompson, 2009). Thus, the differences in neuronal patterns of activation found across conditions in those studies might be due to differences in the visual complexity of the displays used to cue the production of the verb rather than in differences in verb classes and argument structures.

In sum, the brain structures involved in sentence production as well as the factors that affect neural computation during spontaneous sentence production are still understudied. In the current experiment, we address these questions by measuring behavioural and brain responses during sentence production.

Participants produced sentences with different structures (intransitives, transitives, and ditransitives), allowing us to assess the role of verb argument number and thus structural complexity in sentence production. In order to circumvent the problem of using different displays across conditions observed in earlier studies, participants saw identical visual displays with three objects across conditions, thus keeping the total number of nouns to be used in each sentence constant. On each trial, participants first saw a screen with the verb to be used in the upcoming trial (e.g., *“wash”* in the case of the transitive condition), followed by three geometrical objects (a triangle, a circle, and a square), and they were instructed to produce an intransitive, transitive or ditransitive sentence (e.g., *“The triangle, the circle and the square [verb]”; “The triangle and the circle [verb] the square”; “The triangle [verb] the circle to the square”).* Participants were cued to use either existing verbs (i.e., known verbs) or verbs that do not exist in the Dutch lexicon (i.e., pseudo-verbs) in these sentences.

This design allows us to focus on two experimental questions regarding the neurobiology of sentence generation. The *first* experimental question concerns the role of different verb argument structures in modulating production-specific neural responses. We expected to see a graded activation pattern within the language network reflecting the complexity of the sentence structures (in this case, the number of verb arguments): we expected structural complexity to modulate the unification load in the inferior frontal gyrus.

The *second* experimental question concerns the contribution of lexical representations to structural processing during sentence production. We tested whether the neural computation for sentences differs when the lexical entry of the target verb provides syntactic feature information compared to when the verb has no lexical representation (known verbs vs. pseudo-verbs). If argument structure information is verb-bound, there should be more activation during production of sentences using known verbs than pseudo-verbs in regions related to the processing and representation of words within the language network. In contrast, if representations for sentence structures are abstract, there should be no difference between the verb argument structure effect for known verbs and pseudo-verbs beyond activation related to having a memory entry for the known but not the pseudo-verbs.

As we were particularly interested in the brain responses to processing of sentence structure and potential lexical-syntactic components of sentence processing, the analyses focused on two regions of interest linked to syntactic processing: the left inferior frontal gyrus (LIFG) and the left posterior middle temporal gyrus (LpMTG). If the activation of entries in the mental lexicon also entails activation of syntactic information, a graded activation pattern might also be found in the posterior temporal cortex.

## Materials and Methods

### Participants

Thirty right-handed native Dutch students (23 females, *M=21.8* years, range 18-28 years) participated in the experiment in return for course credit or monetary compensation after giving written informed consent. The study was approved by the CMO committee on Research Involving Human Participants (Region Arnhem-Nijmegen). Participants had no history of neurological or language-related disorders, and reported having normal or corrected-to-normal vision and hearing. Two participants were excluded from the analyses: one participant made too many mistakes during the sentence production task (only 69% correct sentence productions), and another participant was excluded because of technical failure during image acquisition, leaving 28 participants for the final analyses.

### Materials

#### Stimuli

Eighteen verbs (six to be used in each construction: intransitive, transitive and ditransitive) were selected and another 18 pseudo-verbs were constructed conforming to Dutch phonotactic rules (also divided into three sets of six pseudo-verbs each). Additionally, three new verbs and three new pseudo-verbs were used as examples for each of the conditions.

Three simple shapes (a square, a circle and a triangle; see Figure 1) were used as stimuli. On all trials, the shapes were presented as white figures on a black background (with a 10-degree visual angle separation), and participants were instructed to produce sentences naming the shapes in a left-to-right order. This presentation eliminated differences in visual input across conditions (and thus eliminated confounds observed in prior studies). The order of the shapes displayed was randomized across trials.

**Figure 1.**
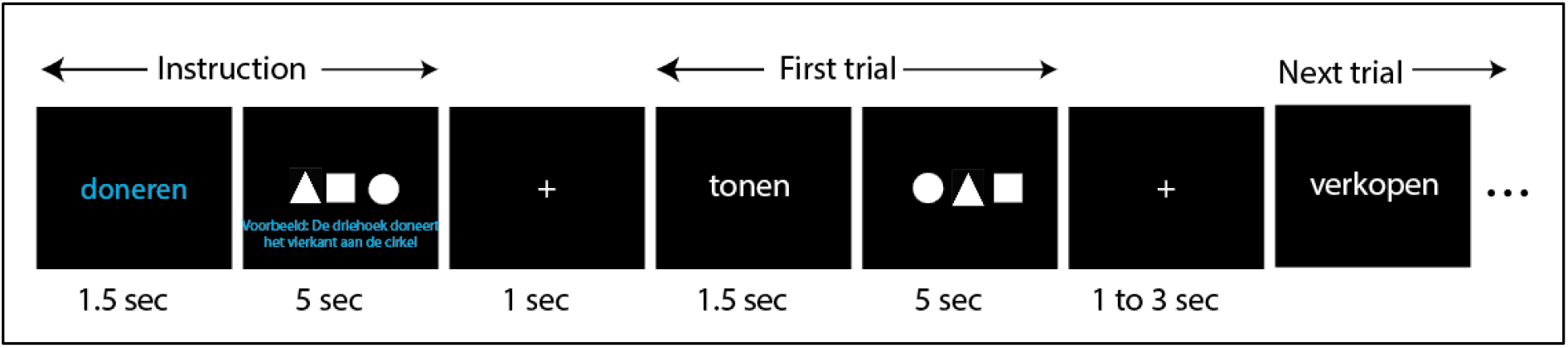
The production task. Each block started with an instruction trial (see “Instruction” example) where the verb was shown in the middle of the screen in blue font for 1.5 sec, followed by three shapes and the required corresponding sentence for 5 sec. Participants were instructed to read the example sentence silently. After a 1 sec interval, the experimental trials started. Each block consisted of six trials and participants were asked to use the same sentence structure as in the example throughout the block. At the beginning of each trial, the (pseudo)verb to be used in that trial was shown in the centre of the screen in white font for 1.5 sec, followed by the three shapes shown for 5 sec. Participants were instructed to overtly produce the sentence while the shapes were on the screen mentioning the three geometrical objects from left to right and using the verb shown prior to picture presentation. The next trial began after a 1-3 sec inter-trial interval. Instruction: *“doneren”* = donate, *“Voorbeeld: De driehoek doneert het vierkant aan de cirkel”* = Example: The triangle donates the square to the circle. Trials: *“tonen”* = show, *“verkopen”* = sell.

### Experimental procedures

#### Sentence production task

Participants were instructed to produce sentences following the example presented on the screen during the instruction phase (Figure 1). Prior to the fMRI session, participants underwent a short practice session outside the scanner to be acquainted with the task using both existing verbs and pseudo-verbs for each of the three verb argument structure conditions. The (pseudo)verbs used in the practice session were not used in the main task.

For the task in the scanner, participants completed 30 blocks of six trials each (for a total of 180 trials for the whole experimental session). Crossing the variable verb argument structure (3 levels) and verb type (2 levels) resulted in six experimental conditions. Conditions were grouped into blocks, so that all sentences within each block belonged to the same condition. As there were 30 blocks, each condition was shown five times. In each block, all six verbs were used only once, but were presented in a different order across blocks.

Each block started with an instruction screen. First, the example verb appeared on screen (visual angle 2.5 to 3.5 degrees) for 1.5 sec, followed by a screen with three shapes (visual angle 10 degrees) and the required sentence structure printed under the shapes for 5 sec. Participants were instructed to read the example silently. They were then given six experimental trials of the same condition (i.e., six trials requiring use of the same sentence structure). In each of these six trials, the (pseudo)verb appeared on screen for 1.5 sec first, followed by three shapes presented for 5 sec. The participants were instructed to produce a sentence with the same sentence structure as in the instruction at the beginning of the block, mentioning the shapes from left to right and using the (pseudo)verb shown just before the presentation of the shapes.

The three sentence structures were: a) intransitives (e.g., “*De driehoek, de cirkel en het vierkant stralen*”; The triangle, the circle and the square shine), b) transitives (e.g., *“De cirkel en het vierkant wassen de driehoek*”; The circle and the square wash the triangle), and c) ditransitives (e.g., “*Het vierkant geeft de driehoek aan de cirkel*”; The square gives the triangle to the circle). Verbs needed to be inflected according to third person singular or plural. Participants were instructed to speak as soon as the three shapes appeared on the screen. The screen was replaced with a central fixation cross after 5 sec and, after a varying inter-trial interval (1, 2, 2.5, or 3 sec), the next trial started. The experiment lasted approximately 30 min.

Participants’ verbal responses were recorded using a noise cancellation system suppressing scanner noise, and analysed offline for accuracy and two production variables: Production Onsets and Production Durations (calculated as the difference between Production Onset and Offset). For the behavioural analyses, we rejected sentences that were unfinished (e.g., *“Het vierkant, de driehoek en de cirkel….”),* used an incorrect structure (e.g., a transitive sentence in a ditransitive block) or an incorrect verb (e.g., *“De driehoek <u>geeft</u> de cirkel aan het vierkant”,* when cued with the verb <*melden*>). Production Onset was calculated as the time between picture onset and start of the utterance, and Production Offset was calculated as the time between picture onset and sentence completion. In case the sentence was not completed within the 5-second trial window, the Production Offset time was coded as unfinished and the trial was rejected. Next, Production Duration was calculated as the time between Production Onset and Production Offset. In Dutch, noun gender is marked with the definite article, with the square (singular form) taking the neuter article “het” *(het vierkant),* and the other two figures with the non-neuter gender article “de” *(de cirkel, de driehoek).* However, because we were interested brain activation areas in production of sentences with correct structures, we included trials where article selection was not always correct (e.g., *de vierkant,* instead of *het vierkant).* All other responses were categorised as incorrect.

#### Overt/covert task

In order to control for brain response related to motion, participants completed another run with ten blocks after the main sentence production task. Here, they were instructed to produce sentences overtly in five blocks and covertly in the remaining five blocks. Each block consisted only of three sentences and only elicited intransitive sentences. The procedure was similar to that of the main task. The two condition blocks appeared in a randomized order. The verb to be used was shown on screen for 1.5 sec, followed by the three figures shown on screen for 5 sec. Participants were instructed to construct a sentence in the same format as the example sentence at the beginning of the block. We then compared BOLD responses to overt and covert sentence production.

### Data acquisition

#### Functional MRI data

MRI data were recorded in a 3T MR scanner (PrismaFit, Siemens Healthcare, Erlangen, Germany) using a 32-channel head coil. Whole-brain functional images were collected using a multi-band (accelerator factor of 8) T2*-weighted sequence: repetition time (TR): 735 ms; echo time (TE) 39 ms; field of view 210 x 210 mm; 64 slices; voxel size 2.4 x 2.4 x 2.4 mm. To correct for distortions, fieldmap images were also recorded.

Additionally, T1-weighted anatomical scans at 1 mm isotropic resolution were acquired with TR 2300 ms, TE 3.03 ms, flip angle 8°, and FOV 256 × 256 × 192 mm.

### Data analysis

#### Behavioural data

Analyses compared accuracy and two production measures (Production Onset and Production Duration) across conditions. Table 1 lists the types and numbers of incorrect responses (incorrect structures, incorrect verb uses, incorrect determiner uses, and combinations of incorrect categories). We compared the number of correctly produced sentences across conditions using mixed-effects logit models (Barr, Levy, Scheepers, & Tily, 2013; Jaeger, 2008; Pinheiro & Bates, 2000) and Production Onsets and Durations using a mixed effects model in R (R Development Core Team, 2012). All three models contained the factors Verb Argument Structure (3 levels: intransitive, transitive, ditransitive) and Lexicality (known verb vs. pseudo-verb). Deviation coding was used for the factor Lexicality and we looked at the linear contrast (first polynomial) for the factor Verb Argument Structure. The models included random effects for participants and items. Following Barr et al. (2013), we report models with the maximal random effect structure leading to convergence. When a model did not converge, we iteratively removed random slopes for factors with the lowest variance one at a time. Consequently, the accuracy model included random slopes for Lexicality for items, while the production onset model included random slopes for Lexicality for participants and the production duration model included random slopes for Lexicality for both participants and items and Verb Argument Structure for participants.

**Table 1.**
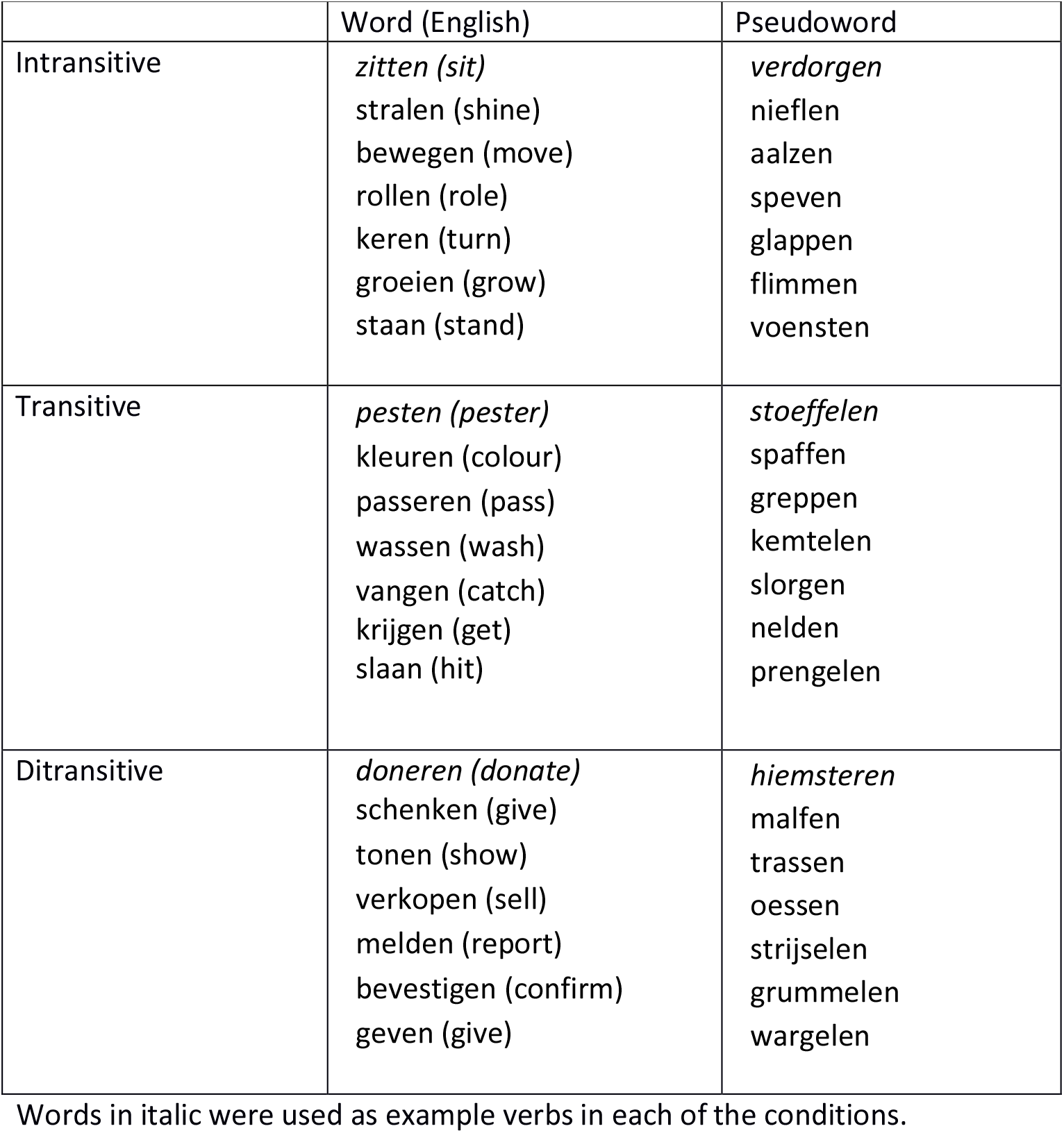
Dutch verbs and pseudo-verbs (with English translations) used in the production task

#### MRI data

First, DICOM images were converted to nifti images. Then functional volumes were realigned using the fieldmap correction using preprocessing tools in SPM12 (www.fil.ion.ucl.ac.uk), and coregistered to the individual structural image and further normalized to a standard MNI space (resampled at voxel size 2 x 2 x 2 mm). Lastly, the images were spatially smoothed with a kernel of 5 mm full width at half maximum.

For the sentence production task, for the first level single subject analysis, we computed a general linear model (GLM) with nine experimental conditions (correct trials for each of the six conditions, all incorrect trials, instruction phase at the beginning of each mini block, and a baseline phase at the offset of the production for all trials), together with six motion parameters and a regressor for each of the affected volumes detected by ArtRepair program (http://cibsr.stanford.edu/tools/human-brain-project/artrepair-software/) as nuisance regressors. For the six experimental condition regressors and the instruction phase regressor, the onset of each trial was defined as the picture onset time, and for the baseline phase, we modelled time just after the production was completed using the Production Offset as the onset of this regressor, convolved with the canonical hemodynamic response function. For each of the experimental conditions, a contrast image of condition minus the baseline phase was computed and compared across participants on the group level.

First, we sought the activity difference between conditions in areas that are known to be involved in syntactic processing. For this, we took functional regions of interest (ROIs) using Neurosynth (http://neurosynth.org/, checked on *21/01/2015).* This programme allowed us to select voxels that are reported to be active in multiple studies relating to a key search word. We selected ROIs using “syntactic” as the key search word, with a threshold of z=6. This revealed two clusters (Figure 3): one in the left inferior frontal gyrus (LIFG) and the other in the left posterior middle temporal gyrus (LpMTG). For these two ROIs, we extracted mean beta values for each of the conditions relative to the baseline phase using MarsBar (http://marsbar.sourceforge.net/) and compared these values using a mixed effects model in R (R Development Core Team, 2012). The model contained the factors Verb Argument Structure (3 levels: intransitive, transitive, ditransitive), Lexicality (known verb vs. pseudo-verb) and Region (LIFG vs. LpMTG). Deviation coding was used for the factors Lexicality and Region and we looked at the linear contrast (first polynomial) for the factor Verb Argument Structure. The model included random effects for participants. To allow convergence, we used the same simplification approach as in the analysis of Production Onsets and Durations; thus the final model included only by-participant random slopes for Region. In the next step, we performed a whole brain analysis using flexible factorial analysis embedded in SPM12, with Lexicality (verb, pseudo-verb) and Verb Argument Structure (intransitive, transitive, ditransitive) as factors of interest. On the group level comparison, we first thresholded the brain responses on the voxel-level at p=.001 uncorrected, and took the cluster-size statistic as the test statistic with Family-wise error corrected *p_FWE_* < 0.05 as the cluster threshold (Hayasaka & Nichols, 2003).

For the Overt/covert task, a similar GLM was run with the conditions Overt and Covert as the main regressors of interest together with example and baseline phases as two extra regressors. Since we cannot derive the Production Offset for the covert condition, we calculated the mean Production Offset time for the intransitive condition in the overt task for each participant, and used this time as a proxy of Production Offset for this run. Furthermore, six motion regressors and affected volume regressors as detected by the ARTrepair programme were included as nuisance regressors to the design matrix. Contrast image overt vs covert was computed for each participant, and this contrast image was tested using a one-sample t-test on the group level. We applied the cluster-level statistic (*p_FWE_* < 0.05) to this comparison as well. Results

### Behavioural results

#### Accuracy

Participants produced more correct sentences in the Ditransitive than the Transitive and Intransitive conditions (mean proportion correct: Ditransitive: .92, Transitive: .81, Intransitive: .79; *β*: .89, z=8.95, *p*<.001) and more correct sentences in the Known verb condition (proportion correct *M*=.91) than the Pseudo-verb condition (*M=.77; β*=-1.24, *z*=-10.6, *p*<.001). The two effects did not interact (|*Z*|<1). In other words, accuracy was highest in the condition where the verb had to be kept in working memory for the shortest amount of time (Ditransitives) and lowest when it had to be kept in working memory the longest (Intransitives). The largest number of incorrect responses in this dataset were due to participants using incorrect verbs (532 responses out of 5040 in total) rather than incorrect structures (78 responses out of 5040 responses in total), and most of these rejections were due to incorrect use of pseudo-verbs (see Table 1 for the distribution of errors across conditions).

**Table 1.** Types of errors aggregated over all participants

| lexical status | verb argument structure | No. of trials | incorrect trials<br>(incorrect verb, construction or object, incomplete utterance) | incorrect construction | incorrect verb | incorrect determiner<br>(trials included in the analysis) |
| --- | --- | --- | --- | --- | --- | --- |
| pseudo verb | intransitive | 840 | 234 | 6 | 173 | 24 |
|  | transitive | 840 | 246 | 35 | 171 | 27 |
|  | ditransitive | 840 | 92 | 18 | 61 | 28 |
| known verb | intransitive | 840 | 118 | 2 | 69 | 28 |
|  | transitive | 840 | 73 | 7 | 37 | 28 |
|  | ditransitive | 840 | 40 | 10 | 21 | 28 |

#### Response times

Production Onsets and Production Durations are shown in Figure 2.

**Figure 2.**
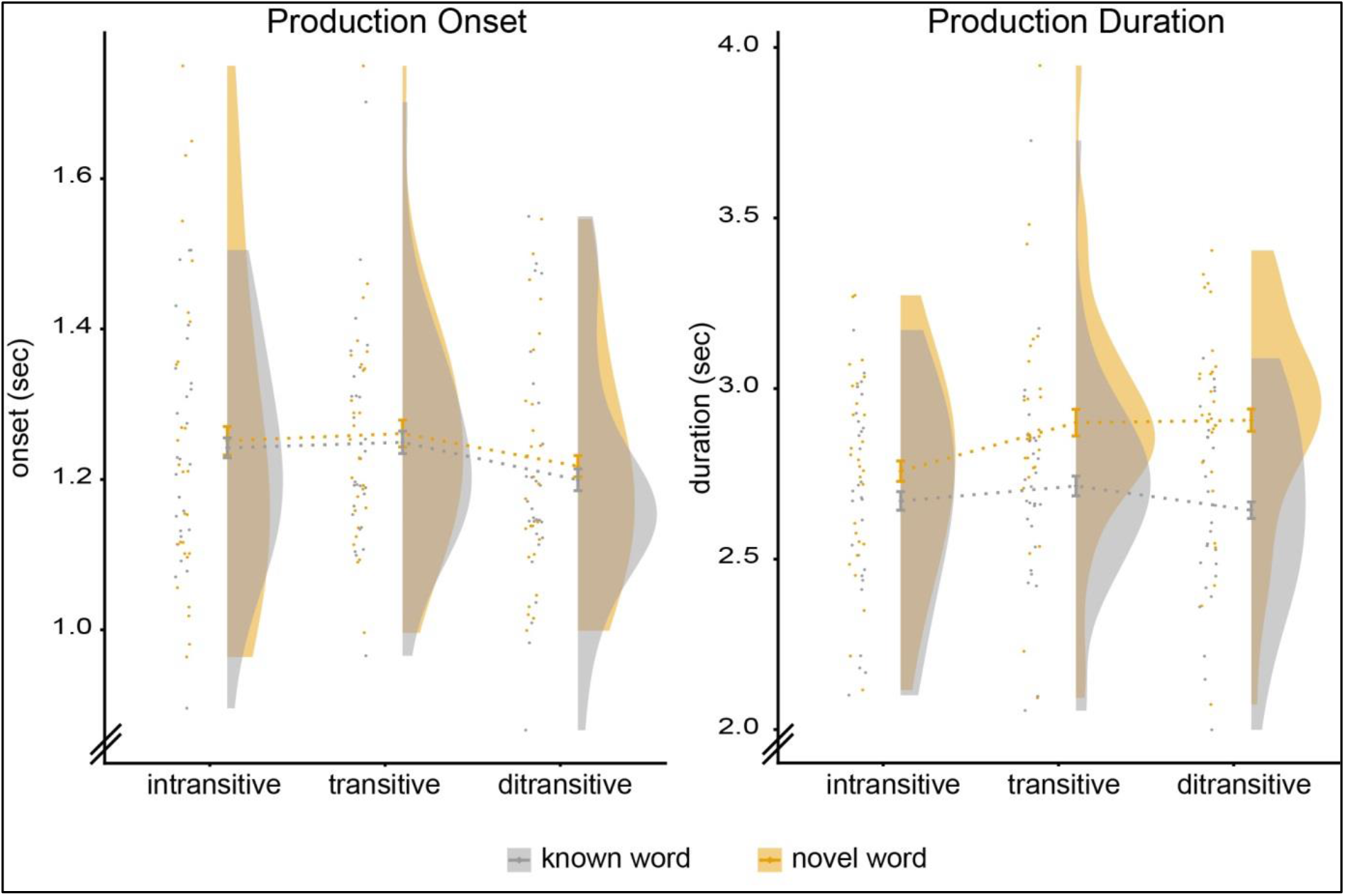
Behavioural results. Left: Production onsets (after picture onset). Right: Production durations (time between production onset and offset). Dots represent individual participants’ data with the mean, standard error of the mean and density represented on their right-hand side.

Production Onsets show how much time speakers needed to prepare the sentence-initial shape name but also vary with structural complexity: sentences beginning with a simple noun phrase are initiated more quickly than sentences beginning with a complex noun phrase (e.g., Smith & Wheeldon, 1999). We performed a mixed effects analysis with fixed effects for Lexicality and Verb Argument Structure, as well as random by-participant and by-item slopes for Lexicality. This analysis showed a main effect of Verb Argument Structure (*β = −0.027, t*=-3.02, *p*=.003). Follow-up comparisons revealed that Ditransitives had shorter onsets than Transitives (*β = .048, t=3.84, p*<.001), while Transitives were not different from Intransitives (*β* =-.01, |*t*|<1). Neither a main effect of Lexicality nor an interaction between Lexicality and Verb Argument Structure were observed (both *t*s<1). This result is consistent with earlier findings showing that speech onsets depend on the phrasal “distance” between the first shape and the second shape to be produced in the sentence, and thus indicate that the timing of sentence production is at least partially subject to structural constraints (e.g., Smith & Wheeldon, 1999).

Production Duration was measured as the time between Production Onset and Production Offset. We performed an analysis with fixed effects for Lexicality and Verb Argument Structure, as well random by-participant slopes for Lexicality and Verb Argument Structure. The results revealed a main effect of Lexicality (*β*= .17, *t*=6.34, *p*<.001), as onsets were faster in the Known verb condition than the Pseudo-verb condition. There was no effect of Verb Argument Structure (*t*=1.68) but there was an interaction between the two factors as the Lexicality effect increased with Verb Argument complexity (*β* = .123, *t*=3.78, *p*<.001). Follow-up tests showed that this effect was driven by an increase in Production Duration over the levels of verb argument for the Pseudo-verbs (*β* =.1, *t*=3.04, *p*=.005), but not the Known verbs (*β* = −.02, |*t*|<1). While we cannot attribute the source of these differences to a specific production process (e.g., word articulation times or the insertion of pauses), the results suggest that producing sentences with pseudo-verbs was cognitively more demanding than producing sentences with known verbs, especially when producing these verbs in a more complex sentence structure.

### Imaging results

#### Sentence production task

##### ROI analyses

First, we considered how the brain areas related to syntactic processing would respond to changes in Verb Argument complexity and Verb lexicality. Our assumption was that there would be a difference in brain responses 1) across sentences with differing numbers of verb arguments (i.e., the Verb Argument Structure effect: Intransitive vs. Transitive vs. Ditransitive), and 2) in sentences using Known words vs. Pseudo-verbs (i.e., the Lexicality effect). For the first point, we expected to find a linear activation increase with an increasing number of verb argument slots to be filled (Intransitive < Transitive < Ditransitive). For the second point, we expected to find differences in brain activation reflecting the presence (Known verb) versus absence (Pseudo-verb) of a lexical memory representation. Moreover, if argument structure information is verb-bound, we should find an interaction with more activation during production of sentences using known verbs than pseudoverbs.

The mixed-effect model on the mean beta values extracted from the two ROIs (LIFG and LpMTG) included the factors ROI (LIFG, LpMTG), Verb Argument Structure (Intransitive, Transitive, Ditransitive), and Lexicality (Known verb, Pseudo-verb). The model revealed a main effect of ROI (*β* =.62, *t*=2.46, *p*=.02), a main effect of Lexicality (*β* =-.19, *t*=-2.78, *p*=.006), and a significant interaction between Lexicality and Verb Argument Structure (*β* =-.3, *t*=-2.53, *p*=.01). No main effect of Verb Argument Structure (linear contrast: *t*<1) nor any additional interaction (all *t*s<|1|) was found

To interpret the interaction of Lexicality and Verb Argument Structure, we ran models per level of the factor Lexicality. These analyses revealed a linear increase with Verb Argument Structure for the Known verbs (*β* =.39, *t*=2.34, *p*=.02), but not the Pseudo-verbs (*β* =-.2, *t*=-1.25, *p*=.2).

##### Flexible factorial: Verb Argument (Intransitive/Transitive/Ditransitive) vs Lexicality (Known verb/Pseudo-verb)

Next, we performed an analysis looking at the whole brain using the flexible factorial analysis with Verb Argument Structure (Intransitive, Transitive, Ditransitive), Lexicality (Known verb, Pseudoverb), and Subjects as factors included in the model.

#### Verb Argument effect

The F-contrast for Verb Argument Structure revealed a large set of areas (see Table 3) with significantly different activation levels among conditions. Of these significant clusters, an increase over Verb Argument Structure levels was found in the LpMTG/AG and the precuneus (see Figure 4A). The rest of the clusters were more active for simpler verb argument structures (intransitive > transitive > ditransitive).

**Figure 3.**
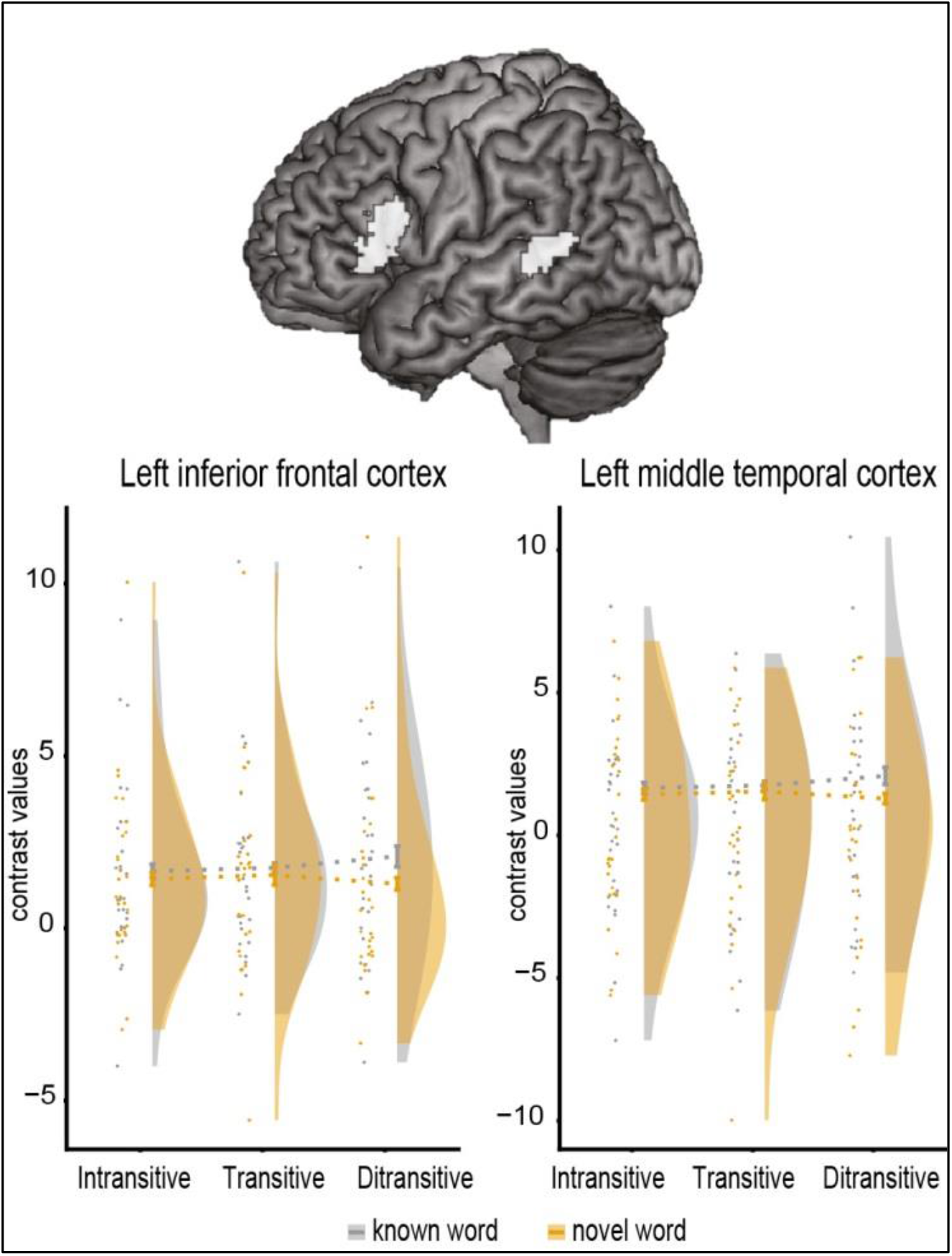
ROI activation pattern. Top: Regions of interest in left inferior frontal (left) and left middle temporal gyrus (right) defined using the term ‘syntactic’ on neurosynth.org. Bottom: Mean beta weights extracted from the predefined region of interest in the left inferior frontal gyrus (left) and the left posterior middle temporal gyrus. Dots represent individual participants’ data with the mean, standard error of the mean and density represented on their right-hand side.

**Figure 4.**
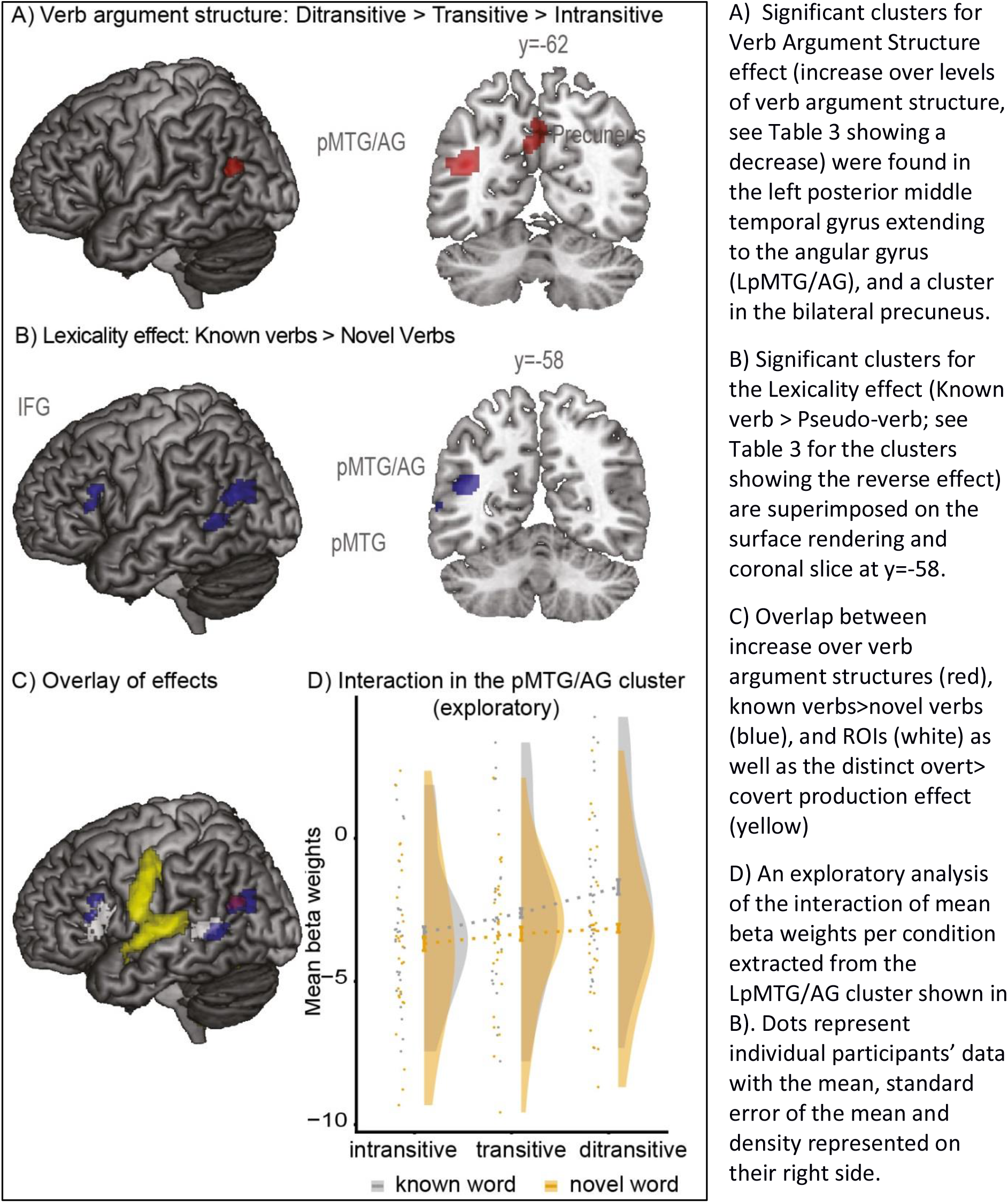
Imaging results. A) Significant clusters for Verb Argument Structure effect (increase over levels of verb argument structure, see Table 3 showing a decrease) were found in the left posterior middle temporal gyrus extending to the angular gyrus (LpMTG/AG), and a cluster in the bilateral precuneus. B) Significant clusters for the Lexicality effect (Known verb > Pseudo-verb; see Table 3 for the clusters showing the reverse effect) are superimposed on the surface rendering and coronal slice at y=-58. C) Overlap between increase over verb argument structures (red), known verbs>novel verbs (blue), and ROIs (white) as well as the distinct overt> covert production effect (yellow) D) An exploratory analysis of the interaction of mean beta weights per condition extracted from the LpMTG/AG cluster shown in B). Dots represent individual participants’ data with the mean, standard error of the mean and density represented on their right side.

**Table 3.**
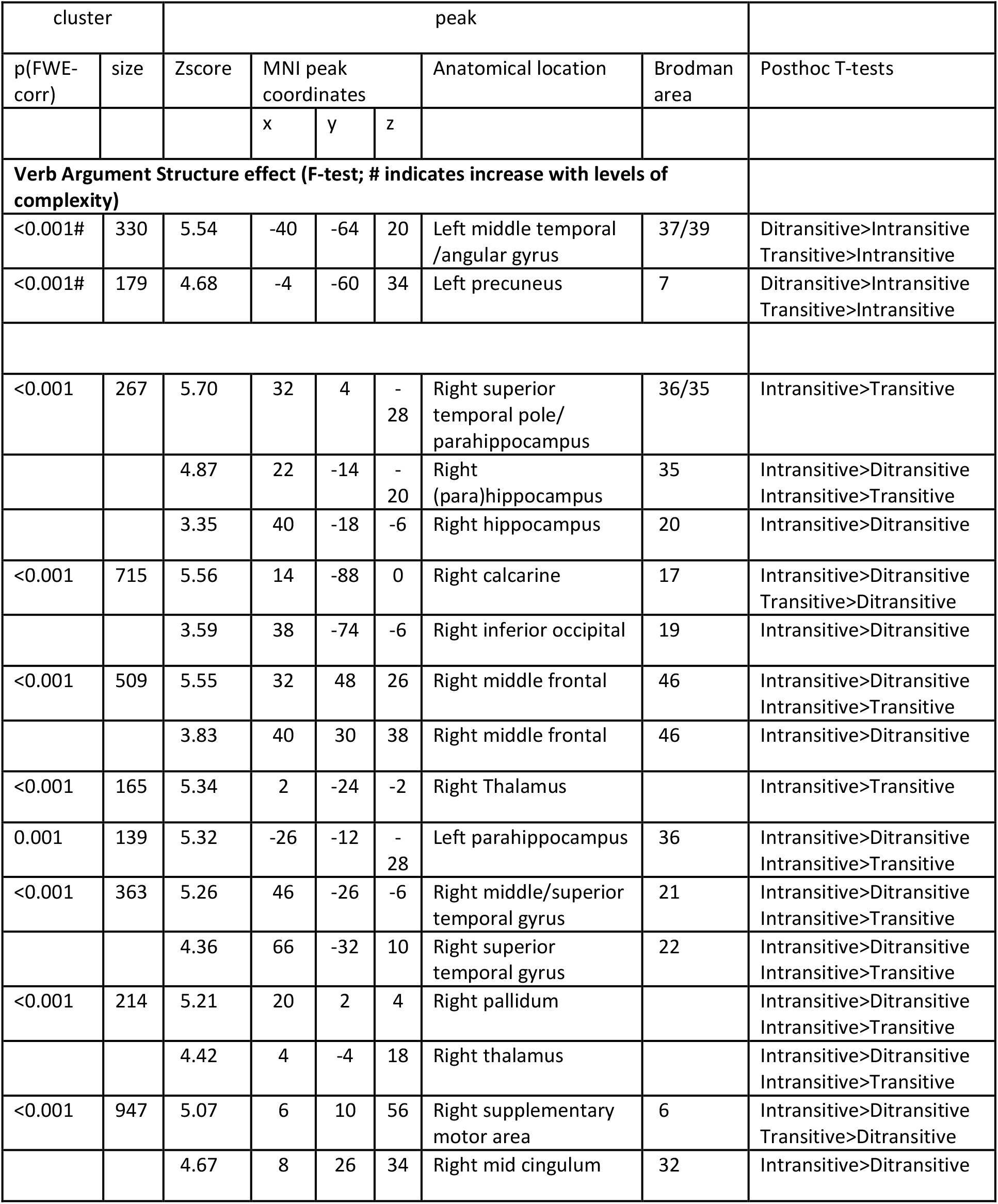
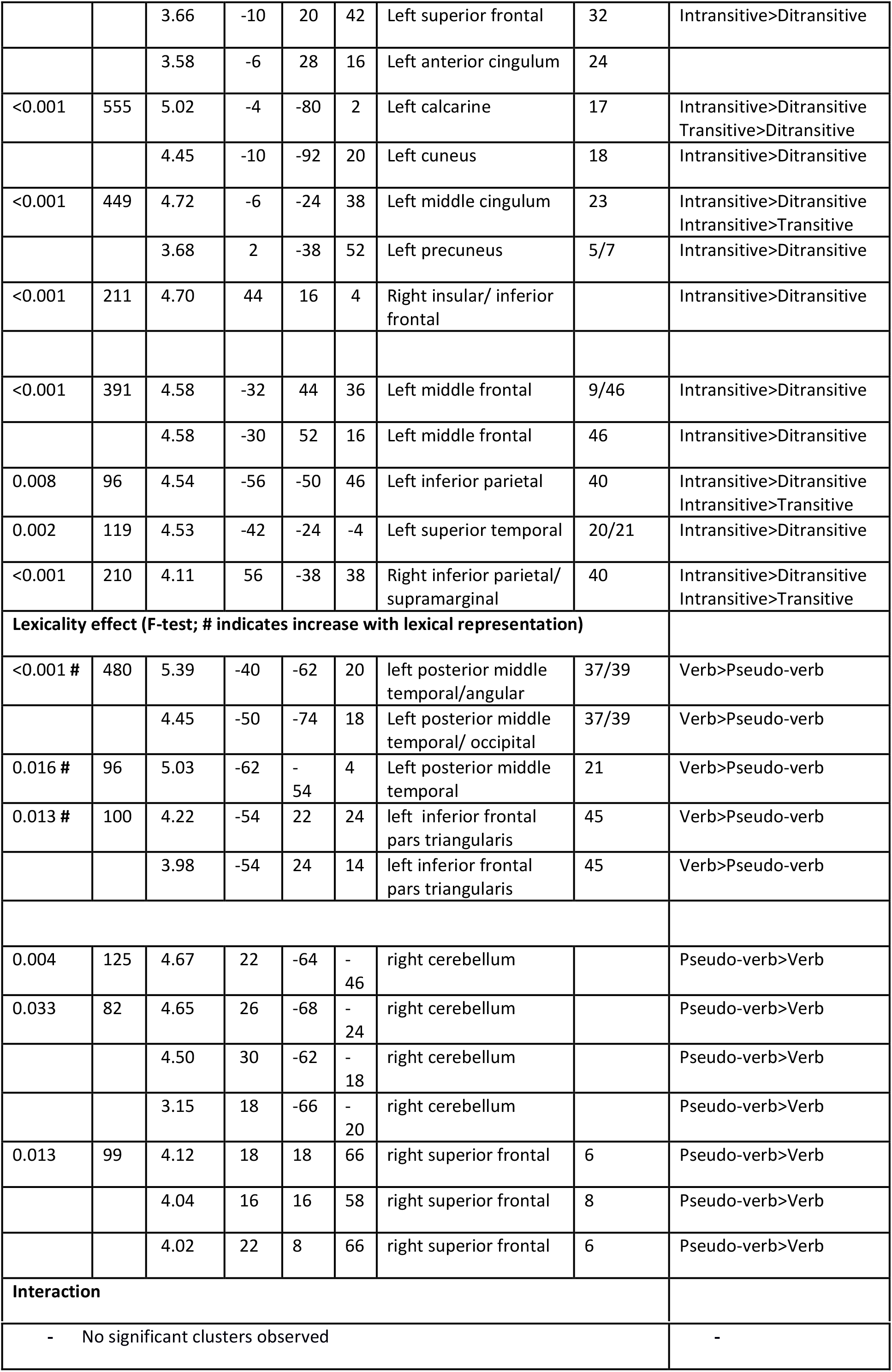
fMRI whole-brain results (local maxima more than 20 mm apart are reported)

| cluster |  | peak |  |  |  |  |  |  |
| --- | --- | --- | --- | --- | --- | --- | --- | --- |
| p(FWE-corr) | size | Zscore | MNI peak coordinates |  |  | Anatomical location | Brodman area | Posthoc T-tests |
|  |  |  | x | y | z |  |  |  |
| **Verb Argument Structure effect (F-test; # indicates increase with levels of complexity)** |  |  |  |  |  |  |  |  |
| <0.001# | 330 | 5.54 | -40 | -64 | 20 | Left middle temporal /angular gyrus | 37/39 | Ditransitive>Intransitive<br>Transitive>Intransitive |
| <0.001# | 179 | 4.68 | -4 | -60 | 34 | Left precuneus | 7 | Ditransitive>Intransitive<br>Transitive>Intransitive |
| <0.001 | 267 | 5.70 | 32 | 4 | -28 | Right superior temporal pole/ parahippocampus | 36/35 | Intransitive>Transitive |
|  |  | 4.87 | 22 | -14 | -20 | Right (para)hippocampus | 35 | Intransitive>Ditransitive<br>Intransitive>Transitive |
|  |  | 3.35 | 40 | -18 | -6 | Right hippocampus | 20 | Intransitive>Ditransitive |
| <0.001 | 715 | 5.56 | 14 | -88 | 0 | Right calcarine | 17 | Intransitive>Ditransitive<br>Transitive>Ditransitive |
|  |  | 3.59 | 38 | -74 | -6 | Right inferior occipital | 19 | Intransitive>Ditransitive |
| <0.001 | 509 | 5.55 | 32 | 48 | 26 | Right middle frontal | 46 | Intransitive>Ditransitive<br>Intransitive>Transitive |
|  |  | 3.83 | 40 | 30 | 38 | Right middle frontal | 46 | Intransitive>Ditransitive |
| <0.001 | 165 | 5.34 | 2 | -24 | -2 | Right Thalamus |  | Intransitive>Transitive |
| 0.001 | 139 | 5.32 | -26 | -12 | -28 | Left parahippocampus | 36 | Intransitive>Ditransitive<br>Intransitive>Transitive |
| <0.001 | 363 | 5.26 | 46 | -26 | -6 | Right middle/superior temporal gyrus | 21 | Intransitive>Ditransitive<br>Intransitive>Transitive |
|  |  | 4.36 | 66 | -32 | 10 | Right superior temporal gyrus | 22 | Intransitive>Ditransitive<br>Intransitive>Transitive |
| <0.001 | 214 | 5.21 | 20 | 2 | 4 | Right pallidum |  | Intransitive>Ditransitive<br>Intransitive>Transitive |
|  |  | 4.42 | 4 | -4 | 18 | Right thalamus |  | Intransitive>Ditransitive<br>Intransitive>Transitive |
| <0.001 | 947 | 5.07 | 6 | 10 | 56 | Right supplementary motor area | 6 | Intransitive>Ditransitive<br>Transitive>Ditransitive |
|  |  | 4.67 | 8 | 26 | 34 | Right mid cingulum | 32 | Intransitive>Ditransitive |
|  |  | 3.66 | -10 | 20 | 42 | Left superior frontal | 32 | Intransitive>Ditransitive |
|  |  | 3.58 | -6 | 28 | 16 | Left anterior cingulum | 24 |  |
| <0.001 | 555 | 5.02 | -4 | -80 | 2 | Left calcarine | 17 | Intransitive>Ditransitive<br>Transitive>Ditransitive |
|  |  | 4.45 | -10 | -92 | 20 | Left cuneus | 18 | Intransitive>Ditransitive |
| <0.001 | 449 | 4.72 | -6 | -24 | 38 | Left middle cingulum | 23 | Intransitive>Ditransitive<br>Intransitive>Transitive |
|  |  | 3.68 | 2 | -38 | 52 | Left precuneus | 5/7 | Intransitive>Ditransitive |
| <0.001 | 211 | 4.70 | 44 | 16 | 4 | Right insular/ inferior frontal |  | Intransitive>Ditransitive |
| <0.001 | 391 | 4.58 | -32 | 44 | 36 | Left middle frontal | 9/46 | Intransitive>Ditransitive |
|  |  | 4.58 | -30 | 52 | 16 | Left middle frontal | 46 | Intransitive>Ditransitive |
| 0.008 | 96 | 4.54 | -56 | -50 | 46 | Left inferior parietal | 40 | Intransitive>Ditransitive<br>Intransitive>Transitive |
| 0.002 | 119 | 4.53 | -42 | -24 | -4 | Left superior temporal | 20/21 | Intransitive>Ditransitive |
| <0.001 | 210 | 4.11 | 56 | -38 | 38 | Right inferior parietal/ supramarginal | 40 | Intransitive>Ditransitive<br>Intransitive>Transitive |
| **Lexicality effect (F-test; # indicates increase with lexical representation)** |  |  |  |  |  |  |  |  |
| <0.001 # | 480 | 5.39 | -40 | -62 | 20 | left posterior middle temporal/angular | 37/39 | Verb>Pseudo-verb |
|  |  | 4.45 | -50 | -74 | 18 | Left posterior middle temporal/ occipital | 37/39 | Verb>Pseudo-verb |
| 0.016 # | 96 | 5.03 | -62 | -54 | 4 | Left posterior middle temporal | 21 | Verb>Pseudo-verb |
| 0.013 # | 100 | 4.22 | -54 | 22 | 24 | left inferior frontal pars triangularis | 45 | Verb>Pseudo-verb |
|  |  | 3.98 | -54 | 24 | 14 | left inferior frontal pars triangularis | 45 | Verb>Pseudo-verb |
| 0.004 | 125 | 4.67 | 22 | -64 | -46 | right cerebellum |  | Pseudo-verb>Verb |
| 0.033 | 82 | 4.65 | 26 | -68 | -24 | right cerebellum |  | Pseudo-verb>Verb |
|  |  | 4.50 | 30 | -62 | -18 | right cerebellum |  | Pseudo-verb>Verb |
|  |  | 3.15 | 18 | -66 | -20 | right cerebellum |  | Pseudo-verb>Verb |
| 0.013 | 99 | 4.12 | 18 | 18 | 66 | right superior frontal | 6 | Pseudo-verb>Verb |
|  |  | 4.04 | 16 | 16 | 58 | right superior frontal | 8 | Pseudo-verb>Verb |
|  |  | 4.02 | 22 | 8 | 66 | right superior frontal | 6 | Pseudo-verb>Verb |
| **Interaction** |  |  |  |  |  |  |  |  |

|  |  |
| --- | --- |
| - No significant clusters observed | - |

Adding Production Onsets into the model (using parametric modulation) did not change the activation pattern in these areas. Thus it is less likely that this contrast reflects brain responses related to shorter reaction time for the Ditransitive condition, leading to higher possibility of mind wandering, which activates the so-called default mode network that includes areas such as the precuneus and the angular gyrus (Raichle & Snyder, 2007).

#### Lexicality effect

An F-contrast for the Lexicality effect (see Table 3) revealed multiple significant clusters. Higher activation was observed for Known verbs than Pseudo-verbs in three clusters: the LpMTG, the LpMTG extending to the angular gyrus (LpMTG/AG), and the LIFG (Figure 4B left). The opposite contrast revealed clusters in the right superior frontal and cerebellar regions.

#### Interaction Lexicality and Verb Argument

At the whole brain level, we did not find any areas showing an interaction between Verb Argument Structure and Lexicality. However, the increase in activation in pMTG/AG cluster overlapped to a great extent with the cluster found for the Lexicality effect (see Figure 4c). To test for interactions between Lexicality and Verb Argument Structure, we carried out a further exploratory analysis within this cluster (mean beta values extracted from the significant cluster of main effect of Verb Argument Structure, see Figure 4D).

This revealed a significant interaction between Lexicality and Verb Argument Structure (linear contrast: *β* =.37, *t*=2.52, *p*=.01). Follow-up tests by level of Lexicality showed an activation increase for known verbs (*β* =1.12, *t*=5.6, *p*<.001), however, only a trend in the same direction was observed for pseudo verbs (*β* =.38, *t*=1.86, *p*=.068)

##### Overt vs Covert production

To verify whether the regions stated above reflected movement-related or motor-related brain activation that may be different across conditions, we contrasted the brain activation pattern observed during overt production to that observed during covert production of intransitive sentences in a similar manner to the main task reported above.

The contrast Overt > Covert was tested using a One-sample t-test on the difference contrast images. This contrast showed activation in bilateral motor areas and superior temporal areas known to process motor output and speech perception, respectively (see Figure 4c yellow; Table 4). The differences between conditions found during the overt production task did not overlap with this contrast apart from the cerebellum, confirming that the cortical areas reported for the production task above are not due to any differences in motor related processes.

**Table 4.** Overt versus covert production fMRI whole-brain results (local maxima more than 20 mm apart are reported)

| cluster |  | peak |  |  |  |  |  |
| --- | --- | --- | --- | --- | --- | --- | --- |
| p(FWE-corr) | size | Zscore | MNI peak coordinates |  |  | Anatomical location | Brodman area |
|  |  |  | x | y | z |  |  |
| **Overt > Covert** |  |  |  |  |  |  |  |
| <.001 | 9436 | 8.71 | -40 | -16 | 36 | Left pre/postcentral gyrus | 3/6 |
|  |  | 7.73 | -16 | -60 | -20 | Left cerebellum |  |
|  |  | 7.39 | 18 | -60 | -20 | Right cerebellum |  |
|  |  | 7.11 | -26 | -12 | -6 | Left putamen |  |
|  |  | 7.02 | -34 | -30 | 16 | Left parietal operculum |  |
| <.001 | 4053 | 7.68 | 48 | -10 | 34 | Right pre/postcentral gyrus | 4 |
|  |  | 6.95 | 40 | -26 | 12 | Right heschl's gyrus |  |
|  |  | 6.67 | 60 | -18 | 4 | Right rolandic operculum | 22 |
|  |  | 5.89 | 64 | -2 | 20 | Right pre/postcentral | 43 |
|  |  | 5.65 | 60 | 2 | -2 | Right planum porale |  |
| <.001 | 200 | 7.24 | 28 | -6 | -8 | Right putamen |  |
| <.001 | 277 | 6.21 | 14 | -20 | 0 | Right thalamus |  |
| 0.002 | 152 | 5.6 | -18 | -28 | 62 | Left pre/postcentral gyrus | 4/6 |
| <.001 | 315 | 5.38 | 20 | 16 | 18 | Right caudate |  |
|  |  | 4.85 | 24 | 38 | 0 | Right middle superior frontal gyrus | 47 |
|  |  | 4.69 | 20 | 38 | 22 | Right middle superior frontal gyrus | 46 |
| 0.002 | 158 | 5.13 | 20 | -26 | 60 | Right pre/postcentral gyrus | 6 |
| <.001 | 314 | 5.11 | -18 | 20 | 20 | Left caudate |  |
|  |  | 3.52 | -22 | 40 | 10 | Left middle superior frontal gyrus | 47 |
| 0.005 | 136 | 5.08 | -18 | 6 | 42 | Left superior frontal gyrus |  |
| 0.03 | 94 | 4.93 | 20 | -46 | 30 | Right posterior cingulate cortex |  |
| <.001 | 1502 | 4.78 | -28 | -64 | 8 | Left calcarine cortex | 19 |
|  |  | 4.57 | 18 | -78 | 16 | Right calcarine cortex | 19 |
|  |  | 4.38 | -16 | -84 | 8 | Left calcarine cortex | 17 |

## Discussion

In this study, participants produced sentences using three different syntactic structures with known verbs and pseudo-verbs. At the behavioural level, the results are consistent with production data reported for sentences with a range of different structures and lexical items: participants were more accurate and faster to produce sentences beginning with simple noun phrases rather than complex noun phrases and sentences using known verbs than pseudo-verbs (Konopka, 2012; Martin, Crowther, Knight, Tamborello II, & Yang, 2010; Smith & Wheeldon, 1999). Thus, despite the use of a simple and repetitive task, we find effects of structural complexity and verb lexicality that reflect differences in processing load normally shown in spontaneous production. More importantly, at the neural level, the Verb Argument Structure effect revealed greater activation with increasing Verb Argument Structure (intransitive < transitive < ditransitive) in the LpMTG cluster extending to the angular gyrus (LpMTG/AG). A Lexicality effect (Known words > Pseudo-words) was found in the LIFG as well as the LpMTG and the angular gyrus. The Verb Argument Structure effect in LIFG, and LpMTG regions was stronger for the Known verb compared to the Pseudo-verb condition, while the LpMTG/AG cluster showed both Verb Argument Structure and Lexicality effects. We discuss each of these findings in turn.

### Verb argument structure effect

Our displays prompted production of sentences with three structures: intransitive, transitive and ditransitive. The difference in structural complexity across these sentences was operationalized as a difference in the number of verb arguments per structure, and this is the main focus of the neuroimaging analyses. However, it is important to note that these structures also differed in verb placement. The verb is sentence-final in intransitive sentences, sentence-medial in transitive and ditransitive sentences, but occurs earlier in ditransitive than transitive sentences. This clearly affected the behavioural results. Participants were more accurate at producing sentences with ditransitive structures than with transitive and intransitive structures, suggesting that early verb placement reduced working memory load as the (pseudo-)verb was not present on the screen at the time of production. Production onsets in Ditransitive sentences were also slightly faster than in the other two sentence types. This is consistent with the observation that planning scope (and thus the onset of articulation) is sensitive to the complexity of the initial noun phrase (e.g., Allum & Wheeldon, 2007; Konopka, 2012; Smith & Wheeldon, 1999): sentences beginning with a simple noun phrase (e.g., *“The circle gives…”)* are normally initiated faster than sentences beginning with a complex noun phrase (e.g., *“The circle and the square wash*…”). The presence of this onset difference in the data suggests that production in this task approximated production in tasks with a less repetitive trial structure and in “natural” production: participants did not wait to begin speaking until they planned an entire sentence, but rather began their sentences as quickly as they could (Levelt, 1989) and encoded the shapes shown on the screen in the order of mention.

By contrast, effects of argument number on production are better captured by neuroimaging data. Looking at the whole-brain imaging results of the correct responses, both a LpMTG/AG region and the precuneus showed greater activation with increasing complexity levels of verb argument structure. In the LpMTG/AG cluster the Verb Argument structure effect was present irrespective of the lexicality of the prompted (pseudo)verbs used, albeit less significant for the pseudo-verb condition. When a sentence requires more argument slots to be filled, compilation/unification of words to be used may demand more computational resources. However, that the effect was stronger in the Known verb condition might reflect the use of the lexical-syntactic entries of specific verbs instead of a less specified abstract template.

Greater activation levels for the (pseudo)verb intransitive and transitive conditions compared to the ditransitive condition in other regions listed in Table 3 including areas such as superior temporal and inferior parietal areas may be driven by the fact that the (pseudo)verb had to be kept in working memory until production. The larger number of incorrect verb responses in the intransitive condition may also arise from failure to maintain the verb in working memory.

Another region that showed the Verb Argument Structure effect (smallest activation for intransitives) was the bilateral precuneus. The precuneus is a higher-order association structure that is known to be involved in multiple processes (Cavanna & Trimble, 2006) including memory retrieval (reviewed in Gilmore, Nelson, & McDermott, 2015) and language processing (reiviewed in Rodd, Vitello, Woollams, & Adank, 2015). Repetition suppression studies have also found this area to be involved in semantic processing (Menenti et al., 2011) and syntactic repetition (Schoot, Menenti, Hagoort, & Segaert, 2014). As this area was not among those in which we expected to show a Verb Argument Structure effect, future studies will have to delineate its functional role.

### Lexicality effect

Overall, participants were able to construct sentences using both known verbs and pseudo-verbs within the appointed time. This finding is consistent with the expectation that speakers have acquired at least partially abstract sentence structure templates in the course of language learning that are not bound to specific verbs (Bock, 1990; Fisher 2000a, 2000b; Frazier, 1987; Konopka & Bock, 2009), and that they are able to apply default rules of verb inflection (Ullman, 2001). Having such template sentence structures (or “abstract” structures not bound to verbs) enables speakers to deduce the word categories of pseudo-words encountered during comprehension (as in the case of the famous poem “Jabberwocky” in Lewis Carroll’s “Through the Looking Glass”) and to use the same sentence template (including the correct verb conjugation) in production, as observed in our study. At the same time, speakers made fewer errors, began production more quickly and completed their sentences faster when using known verbs than pseudo-verbs, suggesting parallel use of the lexical-syntactic information, which is only available in the case of known verbs, when producing sentences.

Considering the imaging results, production of sentences using verbs with a known lexical representation activated areas known for syntactic processing in the LIFG and the LpMTG. Additionally, a more posterior pMTG extending to the angular gyrus (LpMTG/AG) also showed greater activation for the Known verbs relative to Pseudo-verbs. This suggests that production of sentences using known verbs activated the core language network more than when the sentences used verbs without existing lexical representations. These areas very much overlap with those reported in single word comprehension studies (reviewed in Price, 2010), especially for the posterior middle temporal regions that are proposed to store lexical entries. Thus, whether we are comprehending or producing sentences using words we know, processing seems to involve overlapping networks (Hagoort & Indefrey, 2014; Segaert, Kempen, Petersson, & Hagoort, 2013).

The increase in activation in the LIFG that we found for known verbs relative to pseudo-verbs might reflect a process of assemblying the information retrieved from the mental lexicon into a coherent sentence-level representation (Hagoort, 2013). One of the advantages of activating syntactic information in the lexicon by recruiting areas that store mental lexical entries may lead to efficient sentence production, reflected in more accurate and faster responses to known verb trials compared to pseudo-verb trials.

## INTERACTION VERB ARGUMENT STRUCTURE AND LEXICALITY

As mentioned before, the posterior MTG/AG showed a Verb Argument Structure effect that was present for both known and pseudo-verbs but stronger for known verbs, showing an interaction between Verb Argument Structure and Lexicality. The Verb Argument Structure effect for known verbs found in the pMTG/AG cluster is in line with findings from aphasia (Thompson, Bonakdarpour, & Fix, 2010), comprehension (Thompson et al., 2007) and single word production (den Ouden et al., 2009) studies. Lesion in this area causes patients to have difficulty in producing sentences with multiple verb argument structures. Post-stroke treatment targeting the use of sentences with complex argument structure shows changes in this region, accompanied by improvement in behaviour (Thompson et al., 2013). Moreover, the ROIs of the syntax network, LIFG and LpMTG, which is more anterior than the pMTG/AG effect above, also showed an increase over the levels of Verb Argument Stucture for the known verbs that was not present for the pseudo-verbs. These regions constitute the core syntactic processing network (Indefrey & Levelt, 2004; Price, 2012). While our findings did not differentiate between the LIFG and LpMTG, they are nonetheless in line with models that attributed a mental lexicon function to posterior temporal regions and unificiation to LIFG (Hagoort, 2013), as both retrieval of more complex lexical-syntactic representations and their unification might lead to the observed pattern. It remains to be seen whether the more posterior temporal activation extending into the angular gyrus is more related to this lexical-syntactic function or whether its pattern could also be accounted for with a semantic explanation of event information differences related to verb argument structure that has been ascribed to the angular gyrus (Binder, 2016; William Matchin, Liao, Gaston, & Lau, 2019).

### Limitations

As most studies on language production, this study has some limitations. The same verbs were repeated across blocks and the same sentence structure was repeated within each block (although no verbs were repeated within a block). These repetitions might have led to adaptation effects that reduced the magnitude of both the lexicality and verb argument structure effects on production. Specifically, speakers could afford to plan their sentences by encoding shapes one-by-one and from-left-to-right. This is similar to earlier studies requiring sequential object naming (Meyer, Sleiderink, & Levelt, 1998; Meyer, Wheeldon, & Konopka, 2012; Smith & Wheeldon, 1999). This type of radically incremental (object-by-object planning) is advantageous in so far as it minimizes processing load and reduces production costs (Ferreira & Swets, 2002), and may have been further exaggerated by the repetitive nature of the task (Griffin, 2001; Meyer et al., 2012). Furthermore, repeated use of the same structure within each block might have enabled fast reuse of the sentence structure template retrieved in the preceding trial on subsequent trials. Nevertheless, the fact that differences in brain activation were observed across conditions demonstrates the strength of the structural effect. In addition, the number of repetitions per word and the number of sentence structures per block was the same across all conditions, and the order of words within a block as well as condition blocks during the task was randomized across participants; thus, the effects reported here are present over and above the repetition effects.

## Conclusions

Speakers successfully produced sentences with different verb argument structures using both known verbs and pseudo-verbs, although production with known-verbs were faster and more accurate compared to when using the pseudo-verbs. Imaging results revealed that the production process engages the core language network in the left inferior frontal and mid/posterior temporal, as well as the angular cortices. Using a verb that has a mental-lexicon that includes sentence relevant syntactic information may support more effective and efficient production of sentences, especially when the sentence structure is associated with complex verb argument slots.

## Acknowledgements

We would like to thank Maarten van den Heuvel for help with the presentation scripts, Vera van ‘t Hoff in helping with the acquisition and transcription of the data, and Birgit Knudsen for transcription of the data.

## References

Allum, P. H., & Wheeldon, L. R. (2007). Planning scope in spoken sentence production: The role of grammatical units. Journal of Experimental Psychology: Learning, Memory, and Cognition, 33(4), 791.

Barr, D. J., Levy, R., Scheepers, C., & Tily, H. J. (2013). Random effects structure for confirmatory hypothesis testing: Keep it maximal. Journal of Memory and Language, 68(3), 10.1016/j.jml.2012.1011.1001. doi: 10.1016/j.jml.2012.11.001

Barron, H. C., Garvert, M. M., & Behrens, T. E. J. (2016). Repetition suppression: a means to index neural representations using BOLD? Philosophical Transactions of the Royal Society B: Biological Sciences, 371(1705). doi: 10.1098/rstb.2015.0355

Ben-Shachar, M., Palti, D., & Grodzinsky, Y. (2004). Neural correlates of syntactic movement: converging evidence from two fMRI experiments. NeuroImage, 21(4), 1320–1336. doi: https://doi.org/10.1016/j.neuroimage.2003.11.027

Binder, J. R. (2016). In defense of abstract conceptual representations. Psychonomic Bulletin & Review, 23(4), 1096–1108. doi: 10.3758/s13423-015-0909-1

Bock, J. K. (1982). Toward a cognitive psychology of syntax: Information processing contributions to sentence formulation. Psychological Review, 89(1), 1–47. doi: 10.1037/0033-295x.89.1.1

Bock, J. K. (1986). Syntactic persistence in language production. Cognitive Psychology, 18(3), 355–387. doi: https://doi.org/10.1016/0010-0285(86)90004-6

Bresnan, J., Asudeh, A., Toivonen, I., & Wechsler, S. (2015). Lexical-functional syntax (Vol. 16): John Wiley & Sons.

Cavanna, A. E., & Trimble, M. R. (2006). The precuneus: a review of its functional anatomy and behavioural correlates. Brain, 129(3), 564–583. doi: 10.1093/brain/awl004

Chang, F., Dell, G. S., & Bock, K. (2006). Becoming syntactic. Psychological Review, 113(2), 234–272. doi: http://doi.org/10.1037/0033-295x.113.2.234

den Ouden, D.-B., Fix, S., Parrish, T. B., & Thompson, C. K. (2009). Argument structure effects in action verb naming in static and dynamic conditions. Journal of Neurolinguistics, 22(2), 196–215. doi: http://dx.doi.org/10.1016/j.jneuroling.2008.10.004

Ferreira, F., & Swets, B. (2002). How incremental is language production? Evidence from the production of utterances requiring the computation of arithmetic sums. Journal of Memory and Language, 46(1), 57–84.

Frazier, L. (1987). Sentence processing: A tutorial review. Attention and Performance XII. In M. Coltheart (Ed.), The Psychology of Reading (pp. 559–586). London: Lawrence Erlbaum Associates.

Gilmore, A. W., Nelson, S. M., & McDermott, K. B. (2015). A parietal memory network revealed by multiple MRI methods. Trends in Cognitive Sciences, 19(9), 534–543. doi: https://doi.org/10.1016/j.tics.2015.07.004

Griffin, Z. M. (2001). Gaze durations during speech reflect word selection and phonological encoding. Cognition, 82(1), B1–B14. doi: https://doi.org/10.1016/S0010-0277(01)00138-X

Hagoort, P. (2005). On Broca, brain, and binding: a new framework. Trends in Cognitive Sciences, 9(9), 416–423. doi: https://doi.org/10.1016/j.tics.2005.07.004

Hagoort, P. (2013). MUC (Memory, Unification, Control) and beyond. Frontiers in Psychology, 4, 416. doi: 10.3389/fpsyg.2013.00416

Hagoort, P., & Indefrey, P. (2014). The Neurobiology of Language Beyond Single Words. Annual Review of Neuroscience, 37(1), 347–362. doi: 10.1146/annurev-neuro-071013-013847

Haller, S., Radue, E. W., Erb, M., Grodd, W., & Kircher, T. (2005). Overt sentence production in event-related fMRI. Neuropsychologia, 43(5), 807–814. doi: https://doi.org/10.1016/j.neuropsychologia.2004.09.007

Hayasaka, S., & Nichols, T. E. (2003). Validating cluster size inference: random field and permutation methods. NeuroImage, 20(4), 2343–2356.

Indefrey, P., Brown, C. M., Hellwig, F., Amunts, K., Herzog, H., Seitz, R. J., & Hagoort, P. (2001). A neural correlate of syntactic encoding during speech production. Proceedings of the National Academy of Sciences, 98(10), 5933–5936. doi: 10.1073/pnas.101118098

Indefrey, P., & Levelt, W. J. M. (2004). The spatial and temporal signatures of word production components. Cognition, 92(1-2), 101–144. doi: 10.1016/j.cognition.2002.06.001

Jackendoff, R. (2002). Foundations of Language: Brain, Meaning, Grammar, Evolution. New York: Oxford University Press.

Jaeger, T. F. (2008). Categorical Data Analysis: Away from ANOVAs (transformation or not) and towards Logit Mixed Models. Journal of Memory and Language, 59(4), 434–446. doi: 10.1016/j.jml.2007.11.007

Konopka, A. E. (2012). Planning ahead: How recent experience with structures and words changes the scope of linguistic planning. Journal of Memory and Language, 66(1), 143–162.

Malyutina, S., & den Ouden, D.-B. (2017). Task-dependent neural and behavioral effects of verb argument structure features. Brain and Language, 168, 57–72. doi: https://doi.org/10.1016/j.bandl.2017.01.006

Martin, R. C., Crowther, J. E., Knight, M., Tamborello II, F. P., & Yang, C. L. (2010). Planning in sentence production: Evidence for the phrase as a default planning scope. Cognition, 116(2), 177–192.

Matchin, W., & Hickok, G. (in press). The cortical organization of syntax. Cerebral Cortex. doi: https://doi.org/10.31234/osf.io/6394f

Matchin, W., Liao, C.-H., Gaston, P., & Lau, E. (2019). Same words, different structures: An fMRI investigation of argument relations and the angular gyrus. Neuropsychologia, 125, 116–128.

Menenti, L., Gierhan, S. M. E., Segaert, K., & Hagoort, P. (2011). Shared Language: Overlap and Segregation of the Neuronal Infrastructure for Speaking and Listening Revealed by Functional MRI. Psychological Science, 22(9), 1173–1182. doi: 10.1177/0956797611418347

Meyer, A., Sleiderink, A., & Levelt, W. (1998). Viewing and naming objects: Eye movements during noun phrase production. Cognition, 66(2), B25–B33.

Meyer, A., Wheeldon, L., & Konopka, A. E. (2012). Effects of speech rate and practice on the allocation of visual attention in multiple object naming. Frontiers in Psychology, 3, 39.

Pickering, M. J., & Branigan, H. P. (1998). The representation of verbs: Evidence from syntactic priming in language production. Journal of Memory and Language, 39(4), 633–651. doi: http://doi.org/10.1006/jmla.1998.2592

Pinheiro, J., & Bates, D. M. (2000). Mixed-Effects Models in S and S_PLUS. New York: Springer.

Price, C. J. (2010). The anatomy of language: a review of 100 fMRI studies published in 2009. Ann. N. Y. Acad. Sci., 1191, 62–88. doi: 10.1111/j.1749-6632.2010.05444.x

Price, C. J. (2012). A review and synthesis of the first 20 years of PET and fMRI studies of heard speech, spoken language and reading. Neuroimage, 62(2), 816–847. doi: http://dx.doi.org/10.1016/j.neuroimage.2012.04.062

Raichle, M. E., & Snyder, A. Z. (2007). A default mode of brain function: A brief history of an evolving idea. NeuroImage, 37(4), 1083–1090. doi: http://dx.doi.org/10.1016/j.neuroimage.2007.02.041

Rodd, J. M., Vitello, S., Woollams, A. M., & Adank, P. (2015). Localising semantic and syntactic processing in spoken and written language comprehension: An Activation Likelihood Estimation meta-analysis. Brain and Language, 141, 89–102. doi: https://doi.org/10.1016/j.bandl.2014.11.012

Schoot, L., Menenti, L., Hagoort, P., & Segaert, K. (2014). A little more conversation – The influence of communicative context on syntactic priming in brain and behavior. Frontiers in Psychology, 5(208). doi: 10.3389/fpsyg.2014.00208

Segaert, K., Kempen, G., Petersson, K. M., & Hagoort, P. (2013). Syntactic priming and the lexical boost effect during sentence production and sentence comprehension: An fMRI study. Brain and Language, 124(2), 174–183. doi: https://doi.org/10.1016/j.bandl.2012.12.003

Segaert, K., Menenti, L., Weber, K., Petersson, K. M., & Hagoort, P. (2012). Shared Syntax in Language Production and Language Comprehension—An fMRI Study. Cerebral Cortex, 22(7), 1662–1670. doi: 10.1093/cercor/bhr249

Segaert, K., Weber, K., de Lange, F. P., Petersson, K. M., & Hagoort, P. (2013). The suppression of repetition enhancement: A review of fMRI studies. Neuropsychologia, 51(1), 59–66. doi: https://doi.org/10.1016/j.neuropsychologia.2012.11.006

Smith, M., & Wheeldon, L. (1999). High level processing scope in spoken sentence production. Cognition, 73(3), 205–246.

Thompson, C. K., Bonakdarpour, B., Fix, S. C., Blumenfeld, H. K., Parrish, T. B., Gitelman, D. R., & Mesulam, M.-M. (2007). Neural Correlates of Verb Argument Structure Processing. Journal of Cognitive Neuroscience, 19(11), 1753–1767. doi: 10.1162/jocn.2007.19.11.1753

Thompson, C. K., Bonakdarpour, B., & Fix, S. F. (2010). Neural mechanisms of verb argument structure processing in agrammatic aphasic and healthy age-matched listeners. Journal of Cognitive Neuroscience, 22(9), 1993–2011. doi: 10.1162/jocn.2009.21334

Thompson, C. K., Riley, E. A., den Ouden, D.-B., Meltzer-Asscher, A., & Lukic, S. (2013). Training verb argument structure production in agrammatic aphasia: Behavioral and neural recovery patterns. Cortex, 49(9), 2358–2376. doi: https://doi.org/10.1016/j.cortex.2013.02.003

Tomasello, M. (2000). Do young children have adult syntactic competence? Cognition, 74(3), 209–253. doi: https://doi.org/10.1016/S0010-0277(99)00069-4

Tyler, L. K., Marslen-Wilson, W. D., Randall, B., Wright, P., Devereux, B. J., Zhuang, J., … Stamatakis, E. A. (2011). Left inferior frontal cortex and syntax: function, structure and behaviour in patients with left hemisphere damage. Brain, 134(2), 415–431. doi: 10.1093/brain/awq369

Ullman, M. T. (2001). The Declarative/Procedural Model of Lexicon and Grammar. Journal of Psycholinguistic Research, 30(1), 37–69. doi: 10.1023/a:1005204207369

Vosse, T., & Kempen, G. (2000). Syntactic structure assembly in human parsing: a computational model based on competitive inhibition and a lexicalist grammar. Cognition, 75(2), 105–143. doi: http://dx.doi.org/10.1016/S0010-0277(00)00063-9

